# Characterizing the tumor microenvironment of metastatic ovarian cancer by single cell transcriptomics

**DOI:** 10.1101/2020.02.11.944561

**Authors:** Susan Olalekan, Bingqing Xie, Rebecca Back, Heather Eckart, Anindita Basu

**Affiliations:** Department of Medicine, University of Chicago; Center for Nanoscale Materials, Argonne National Laboratory

**Keywords:** Ovarian cancer, scRNA-seq, transcriptomics, immune cells, human cancer

## Abstract

Epithelial ovarian cancer is a highly heterogenous, metastatic and lethal disease. The presence of CD8^+^T cells within ovarian tumors is positively associated with overall patient survival. Determining if a patient has T cells that respond to immunotherapies, their characteristics and how they can be manipulated to target cancer cells is an area of intense investigation in cancer therapy. This study determines the cellular composition and the transcriptional state of immune cells in metastatic ovarian cancer samples from patients using single cell RNA sequencing (scRNA-seq). Hierarchical clustering stratified our patient cohort into 2 main groups: 1) a high T cell infiltration (high T_inf_) group and 2) a low T cell infiltration (low T_inf_) group. To assess the immune response in these patient samples, we performed an unsupervised clustering of the T cell population in each group. The T cell population clustered into 4 and 3 subpopulations in the high T_inf_ T cell and low T_inf_ respectively. A granulysin expressing T cell cluster was identified and unique to the High T_inf_ group. Interestingly, although both groups had resident memory CD8^+^T (CD8^+^Trm) cells, only the CD8^+^Trm cells in the high T_inf_ group expressed TOX, a recently described transcription factor. TOX confers longevity to T cells within immunosuppressive environment such as cancer. Interestingly, along with TOX^+^ T cells, we found a unique plasmablast cluster and an IRF8^+^ macrophage cluster unique to the high T_inf_ group. Our comprehensive scRNA-seq study provides important insights in elucidating the immune response in ovarian cancer patients.

## Introduction

Ovarian cancer is the most lethal malignancy of the female reproductive tract^1^. Conventional therapy involving cytoreductive surgery and chemotherapy are 90% effective when cancer is diagnosed at the early stage when it is still restricted to one or both ovaries. Unfortunately, most ovarian cancer cases are diagnosed at stage III or IV when the cancer has metastasized and the diagnosis in these patients result in a 30% 5-year survival rate^2^. To develop efficacious therapies for metastatic ovarian cancer, we need to define the cellular heterogeneity and the transcriptional state within the tumor microenvironment. Immunohistochemical staining and flow cytometry have been useful in categorizing the cell types based on specific cell surface markers but mask intra-cellular heterogeneity. Bulk RNA profiling has been used to categorize high grade serous ovarian carcinoma (HGSOC), the most common and lethal histotype of ovarian cancer, into molecular subtypes^3,4^. However, bulk RNA sequencing averages gene expression and fails to identify the respective contribution of cell subsets. Alternatively, single cell RNA sequencing (scRNA-seq) has emerged as a powerful tool to interrogate tumor composition, revealing previously uncharacterized cellular heterogeneity and gene regulatory networks at single cell resolution^5–10^. Recently, a scRNA-seq study, investigated the heterogeneity in the proposed cell of origin of HGSOC and revealed a high epithelial-mesenchymal transition (EMT) high subtype associated with poor prognosis^11^. These transcriptomic studies have focused on ovarian cancer cells; however, the tumor microenvironment is made up of other cell types that can affect patient stratification, targeted treatment and outcome.

The omentum is mainly an adipose tissue in the peritoneal cavity containing aggregates of immune cells in areas called milky spots^12^. These milky spots act similar to lymph nodes, collecting and responding to antigen within the peritoneal cavity. Interestingly, ovarian cancer cells preferentially colonize adipose tissue with milky spots in the peritoneal cavity^13^. Additionally, adipocytes provide adipokines and act as a source of energy for ovarian cancer cells^14^. These factors prime the omentum as a premetastatic niche for ovarian cancer. The initial presence of ovarian cancer cells in the omentum leads to a recruitment of macrophages into the milky spots without anti-tumor effects. Contrastingly, the presence of tumor infiltrating CD8^+^ T cells in both the ovary and omentum is associated with significantly longer overall survival^15^. However, checkpoint inhibitors, a cancer immunotherapeutic approach, aimed at restoring CD8^+^ T cell function have had a low response rate^15^. We therefore need a better understanding of the tumor microenvironment to improve patient response rate to cancer immunotherapy.

In this current study, we use Drop-seq technology to examine the cells within omental tumors from 6 patients with different histotypes of ovarian cancer. We identified 12 cell clusters and patient samples were stratified into two groups based on immune signatures: 1) a high T cell infiltration (high T_inf_) group and 2) a low T cell infiltration (low T_inf_) group. This unprecedented concurrent transcriptomic analysis of all cells within metastatic ovarian cancer unravels patient-specific responses to cancer and identifies patients that might be suitable for immune checkpoint inhibition.

## Materials and methods

### Tissue collection

Ovarian cancer tissue was collected from women undergoing debulking surgery at the University of Chicago. Human tissue acquisition after patient deidentification was approved by the university of Chicago institutional review board for human research. Ovarian cancer tissue was histologically carcinoma classified and staged by a pathologist according to tumor-node-metastasis (TMN) and/or international federation of gynecology and obstetrics (FIGO) classifications (Table 1). Three of the ovarian cancer samples originated from the left fallopian tube with associated serous tubal intraepithelial carcinoma (STIC).

**Table 1:**
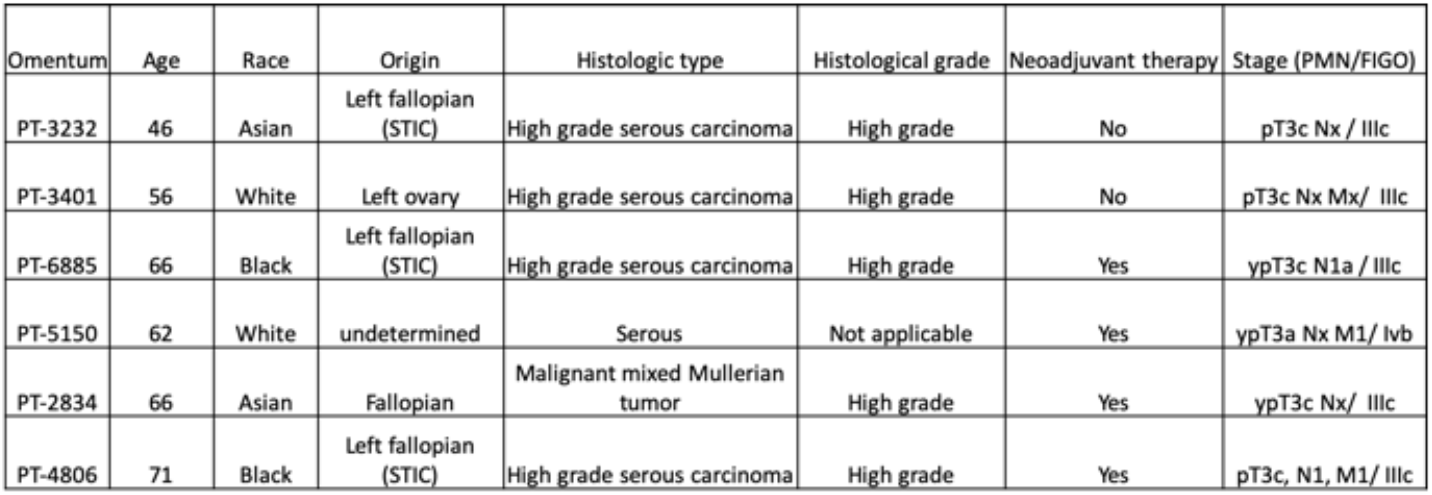
De-identified meta-data for all omental tumors collected from ovarian cancer patients.

### Immunohistochemistry

Ovarian cancer tissues obtained from the anatomic pathology department were fixed overnight in 4% formaldehyde at 4°C. After serial dehydration, the tissues were embedded in paraffin and cut into 5 μm thick sections. Histological evaluation was done with hematoxylin and eosin (H&E). Immunohistochemical staining was performed to confirm the presence of cytokeratin-7 (Thermofisher), pan-vimentin (DAKO), CD45 (Agilent), CD3 (Human Envision) and CD19 (Abcam) cells. Briefly, sections were deparaffinized and rehydrated using xylene and serial dilutions of EtOH in distilled water. Tissue sections were incubated in citrate buffer, pH 6 and heated in a microwave oven. Anti-cytokeratin-7 (1:1000), anti-vimentin (1:100), anti-CD45 (1:100), anti-CD3 (1:100), and anti-CD19 (1:200) antibodies were applied on tissue sections with one-hour incubation at room temperature in a humidity chamber. The antigen-antibody bindings were detected with labeled polymer-HRP Envision system (DAKO, K4007) and DAB+ chromogen (DAKO, K3468) system. Tissue sections were briefly immersed in hematoxylin for counterstaining and were covered with cover glasses.

### Tissue digestion, red blood cell lysis and dead cell removal

Ovarian cancer tissue was transported in DMEM/F12 containing 10% FBS and 1% P/S (10% DMEMF/12) on ice from the anatomic pathology department to the laboratory. The tissue was minced manually with a scalpel and enzymatically digested using 1.5 mg/ml collagenase IV (Sigma-Aldrich), 1 mg/ml hyaluronidase (Sigma-Aldrich) and 500 μg/ml DNase I (GoldBio) in Hank’s balanced salt solution (HBSS) in a 37 °C shaker (200 rpm) for 0.5 – 2 h. Following digestion, cells were resuspended in 10% DMEMF/12 and filtered serially through 70 μm and 40 μm strainers. Red blood cells were lysed by incubating the cell suspension in RBC lysis buffer (Sigma-Aldrich) for 2-5 minutes. Lysis was quenched by adding excess 10% DMEMF/12. The number of live cells was enriched using the dead cell removal kit (Miltenyi, 130-090-101).

### Drop-seq experiments

Drop-seq experiment was performed as previously described^16^. Briefly, cells and barcoded beads were loaded at a concentration of 100,000 cells/ml in PBS-BSA and 120,000 beads/ml in Drop-seq lysis buffer in 3 ml syringes. Droplets were generated using a 125-micron microfluidic device at 16 ml/hr (oil), 4 ml/hr (cells) and 4 ml/hr (beads) with ~15 minutes per collection. Following collection, drops were broken and barcoded beads with mRNA hybridized onto them were collected and washed. Barcoded cDNA attached to the beads or STAMPS were generated by reverse transcription, treated with exonuclease 1 and the number of STAMPs was counted. 5000 STAMPSs were aliquoted per well in a 96-well plate and the cDNA attached to the STAMPS were amplified through 14 PCR cycles. Supernatants from each well were pooled and cleaned with Ampure beads. Purified cDNA was quantified using Qubit 3.0 (Invitrogen) and 450-650 pg of each sample was used as input for Nextera reactions (12 cycles). Tagmented libraries were quantified using Agilent BioAnalyzer High sensitivity chip before submission for sequencing on Illumina’s NextSeq 500, using 75 cycle v3 kits. Paired end sequencing was performed with 20 bp for Read 1 and 64 bp for Read2 using Read1 primer, GCCTGTCCGCGGAAGCAGTGGTATCAACGCAGAGTAC and 5% Illumina PhiX Control v3.

### Data processing, alignment and clustering analysis

From alignment of the reads from the Dropseq experiments, read count matrices for unique molecular identifiers (UMIs) were generated for both exon and intron regions in the human genome (gencode hg38 v.27) using the STAR version 2.5.3 aligner^17,18^. The ovarian cancer samples from six patients’ metastatic omentum were sequenced with Drop-seq. A total of 13 sequencing runs were performed for six Drop-seq samples where each sample was sequenced at least twice to maximize the sequencing depth (PT-3401 was sequenced three times). Each run produced paired-end reads, with Read 1 representing the 12 bp cell barcode and six bp unique molecular identifier (UMI), and Read 2 representing a 60 bp mRNA fragment. Paired-end reads from the same samples were merged to generate six paired-end fastq files. A *snakemake* pipeline was applied to each sequencing run and produce a count matrix representing the gene expression in every cell^19^. Individual count matrices were produced for each of the six patients after accounting for UMI duplicates.

The individual count matrix was produced for each patient with a total number of six count matrices. The summarized counts for each gene were inferred based on both exon and intron reads to produce the gene expression matrix per sample. Expression matrices of six ovarian cancer samples from the metastatic sites in the omentum were generated. To select high quality cells, we applied a filter based on the number of genes detected per cell. Prior to filtering, each sample produced approximately 5000 cells. Based on the median number of genes captured, cells with less than 800 genes were removed from the data sets. For lower quality samples, the threshold was lowered to 600 or 700 as shown in Table 2. We followed a standardized pipeline using single cell analysis tool suite, Seurat v3.0.2^20,21^. A global-scaling logarithmic normalization method was applied to all samples^21^ that normalizes the feature expression counts for each cell by the total expression counts, multiplies by a scale factor of 10,000 (TP10K), and natural-log transforms, as ln(TP10K+1). Each normalization matrix was then scaled by a linear transformation to center the mean gene expression of cells and normalized the gene expression variance. The most variable genes were extracted for principal component analysis (PCA). We applied PCA on the normalized expression matrix with the most variable genes to extract the top 50 components in the data, followed by a heuristic elbow plot on the variance explained of each PC. We selected the number of top variant PCs based on the elbow plot which varies from 10 to 20 depending on the sample. The top PCs were used in further exploration of the data, such as UMAP /tSNE dimension reduction, construction of K-nearest neighbor graph, etc. For analysis that included multiple samples, integration through anchoring^22^ was applied. Note that a subset of genes, usually highly variable ones, was selected to perform the integration, where the integrated gene expression matrix has a lesser number of features (genes) than the original gene expression matrix. The samples from multiple patients were integrated before classifying cell types. Differential expression analysis was done through *FindMarkers* function in Seurat V3 using the Wilcoxon Rank Sum test, and statistically significant markers were extracted for sub-populations or contrast groups based on an adjusted p-value threshold of 0.05. We used dimension reduction methods, UMAP^23^ and tSNE^24^ to generate 2D plots to visualize different cell populations in the experiment.

**Table 2.** Filter parameters applied to samples collected from six patients and the resulting number of cells and genes per sample.

|  | Gene Cutoff | No of Genes | No of Cells |
| --- | --- | --- | --- |
| PT-5150 | 600 | 24776 | 1244 |
| PT-3401 | 800 | 32024 | 3451 |
| PT-6885 | 800 | 24063 | 1071 |
| PT-3232 | 700 | 19297 | 1102 |
| PT-4806 | 700 | 26499 | 1108 |
| PT-2834 | 800 | 26221 | 1909 |

### Cancer subtype classification and correlation with The Cancer Genome Atlas

Four cancer subtypes-differentiated, immunoreactive, mesenchymal, and proliferative were determined from bulk sequencing study on ovarian cancer^4^. Modular scores^25^ were generated between gene expression levels for each cell with upregulated marker signatures on the four subtypes. The subtypes were then assigned to individual cells by the highest positive modular score and if no positive modular scores were found, the subtype was considered undecided.

### Cell type classification and correlation with CellAtlas

To assign cell types to individual cells, we used a bulk RNA sequencing data-set from 95 cell lines collected by CellAtlas^26^, covering 33 major cell types in normal human tissue, including common immune, endothelial, epithelial, fibroblast, mesodermal cells and so on. The cell lines can be further divided into 60 subtypes. We inferred the similarity between individual cells in the samples and cell lines by calculating the pairwise Pearson correlation matrix *C* = {*cor*(*i,j*)} among any cell *i* from the Drop-seq experiments with any cell line, *j* in the CellAtlas. A shared nearest neighbor (SNN) modularity optimization-based clustering^27^ on the CellAtlas cell lines grouped the cell-lines into larger and more general cell-type categories. This was done to eliminate any bias in CellAtlas dataset and remove rare or under-powered cell-types. The resulting clusters of cell-lines were annotated by the most frequent major cell-type from the cell-line groups. For each cell in our cancer sample, cell-type was then assigned from top 5 highly correlated major cell-type clusters. We collapsed ambiguous cell types with neighboring cell types, based on expression profiles.

### Classification with cluster markers, canonical genes, and genetic functions

Sometimes the Gene Ontology (GO) and Pathway analysis gives general/noisy functional categories which make certain cell types hard to identify. By leveraging prior knowledge from the CellAtlas mapping, we were able to locate the relevant function categories for those cell types and narrow down their marker genes efficiently^26^. Moreover, by the consensus of both GO and CellAtlas analyses, we obtained higher confidence in classifying cell types in each patient sample.

We used the cell types obtained from CellAtlas correlation as baseline and curated cell-types manually using canonical genes and functional association. Differentially expressed marker genes were extracted from cell clusters. Cross-referencing was done between gene markers and known canonical genes to the cell-types. Gene ontology and pathway enrichment was performed on gene markers to provide additional evidence for cell-type assignment. We identified 9 major cell types including epithelial cells, fibroblast cells, three immune cell types; B cells/B cell plasma, T cells, and monocytes, mesenchymal stem cells (MSC), embryonic stem cells (ESC), and endothelial cells.

## Results

### Characteristics of the tumor microenvironment

Ovarian cancer samples (1-3 different sections) were collected from the omental metastatic site of six patients (Table 1). Four patients were diagnosed with advanced high-grade serous cancer, one with serous carcinoma and one with malignant mixed Mullerian tumor (MMMT). Ages of the patients ranged from 46 to 71 years. Four of the patients received neo-adjuvant therapy prior to surgery, including the patient diagnosed with MMMT. Normal omentum is comprised mainly of adipocytes as well as aggregates of leukocytes called milky spots^12^. As metastasis progresses, there is an inverse relationship between ovarian tumor growth (cancer and cancer associated cell growth) and adipocytes within the omentum^14^. Therefore, the area of tumor occupied by adipocyte decreases with disease score^28^. To quantify the transformation of the omentum by ovarian cancer cells, we used Imagescope, a digital histopathology software, to annotate and calculate the area occupied by adipocytes in each patient sample on hematoxylin and eosin (H&E) stained sections (Figure 1A). The area of adipocytes was reported as a percentage of the total surface area of the sample and used as a measure of disease score (Figure 1B), similar to Pearce et al.^28^ We rated the patient samples from lowest to highest disease score; PT-5150, PT-3401, PT-6885, PT-3232, PT-4806, PT-2834 respectively. The percentage of cancer cells, stromal cells and immune cells were quantified by immunohistochemistry (IHC) using antibodies against cytokeratin-7 (CK-7), vimentin and CD45 antibodies (Figure 1C-H). There was a varying proportion of each cell type across patients. Notably, we observed that malignant cells in some patients (PT-5150, PT-3232, PT-4806 and PT-2834) double stained for CK-7 and vimentin, suggesting active epithelial-mesenchymal-transition (Figure 1C-F). In addition, the MMMT (PT-2834) sample had two different types of malignant cells that stained separately for CK-7 and vimentin. We assessed the relationship between the area of adipocytes and the cancer and stromal compartments and only observed a positive correlation with CD45^+^ immune cells (Supplementary Figure 1). Notably, we observed aggregation of immune cells closer to adipocytes and sparsely otherwise throughout the rest of the tissue. Together, these data reveal patient variability in the cellular transformation of tumors from the same pathological stage.

**Figure 1.**
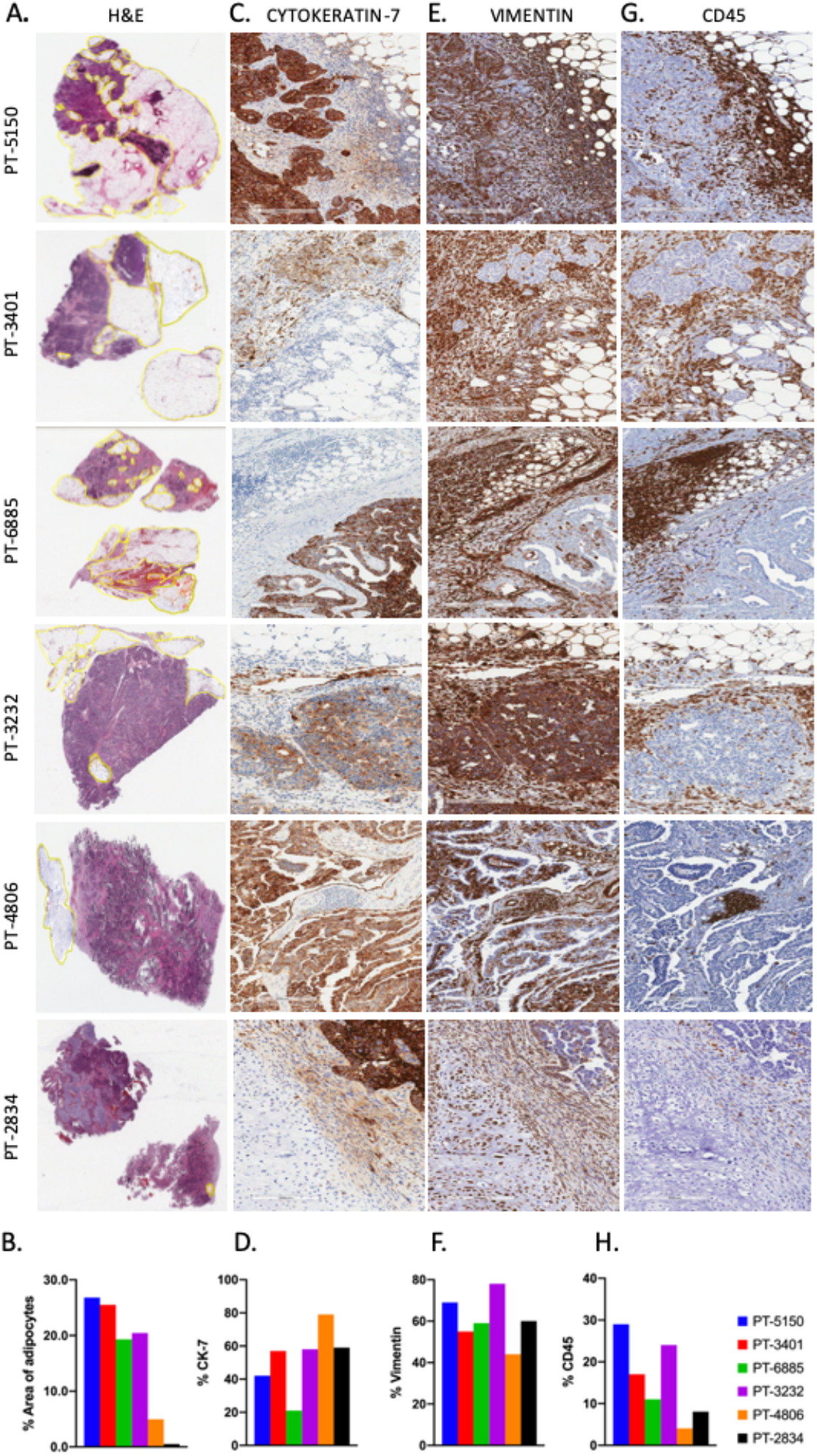
Immunohistochemical staining and sample description. A) The hematoxylin and eosin (H&E) stained sections with B) the histogram of percentage area of adipocytes on the patient sample. C) Cytokeratin-7 (CK-7) staining on patient sample with D) histogram of the percentage of CK-7 positive cells. E) Vimentin staining on patient sample with F) the histogram of percentage of Vimentin positive cells. G) CD45 staining on patient sample with H) histogram of the percentage of CD45 positive cells. Images were taken at A) 40x; C), E), G) 400x.

### Generation of single cell data, cell clustering and cell type assignment

For Drop-seq experiments, tumors samples were enzymatically digested into a single cell suspension and processed^16^. We integrated all six omental samples into a gene expression matrix which contains the expression values of 9885 cells and 40947 genetic features such as protein coding genes, pseudogenes, lncRNA, mapped from the GENCODE human reference genome (version GRCh38). A minimum cut off of 600-800 genes per cell was set, as shown in Table 2. Hierarchical clustering was performed using resolution of 0.2 with 12 clusters detected (Supplementary Figure 2A). Each cluster had cells from all samples (Supplementary Fig. 2B). The cell types were assigned and curated by cell line correlation, canonical genes and functional categories for the significantly differentially expressed genes on the detected clusters. 9 major clusters out of 12 were identified with cell types (Figure 2A). This included 1 cluster for malignant epithelial cells, 1 cluster for fibroblasts, 1 cluster for mesenchymal stem cells, 1 cluster for embryonic stem cells, 1 cluster for endothelial cells. Each patient sample had a varying proportion of the 9 identified cell types/clusters (Figure 2B). Previous bulk mRNA and microRNA expression studies have established 4 transcriptional subtypes of HGSOC^3,29^. To assign molecular subtypes, we extracted gene sets defining each molecular subtype and applied them to our Drop-seq data. At the single cell level, each molecular subtype is represented in every patient tumor sample (Figure 2C). We further analyzed which cell type/cluster belonged to each molecular subtype. The differentiated and proliferative subtypes were mainly made up of epithelial cells, mesenchymal and immunoreactive subtypes were mainly made up of fibroblasts and immune cells respectively suggesting that TCGA classification is reflective of the proportion of cells types within each tumor (Figure 2D).

**Figure 2.**
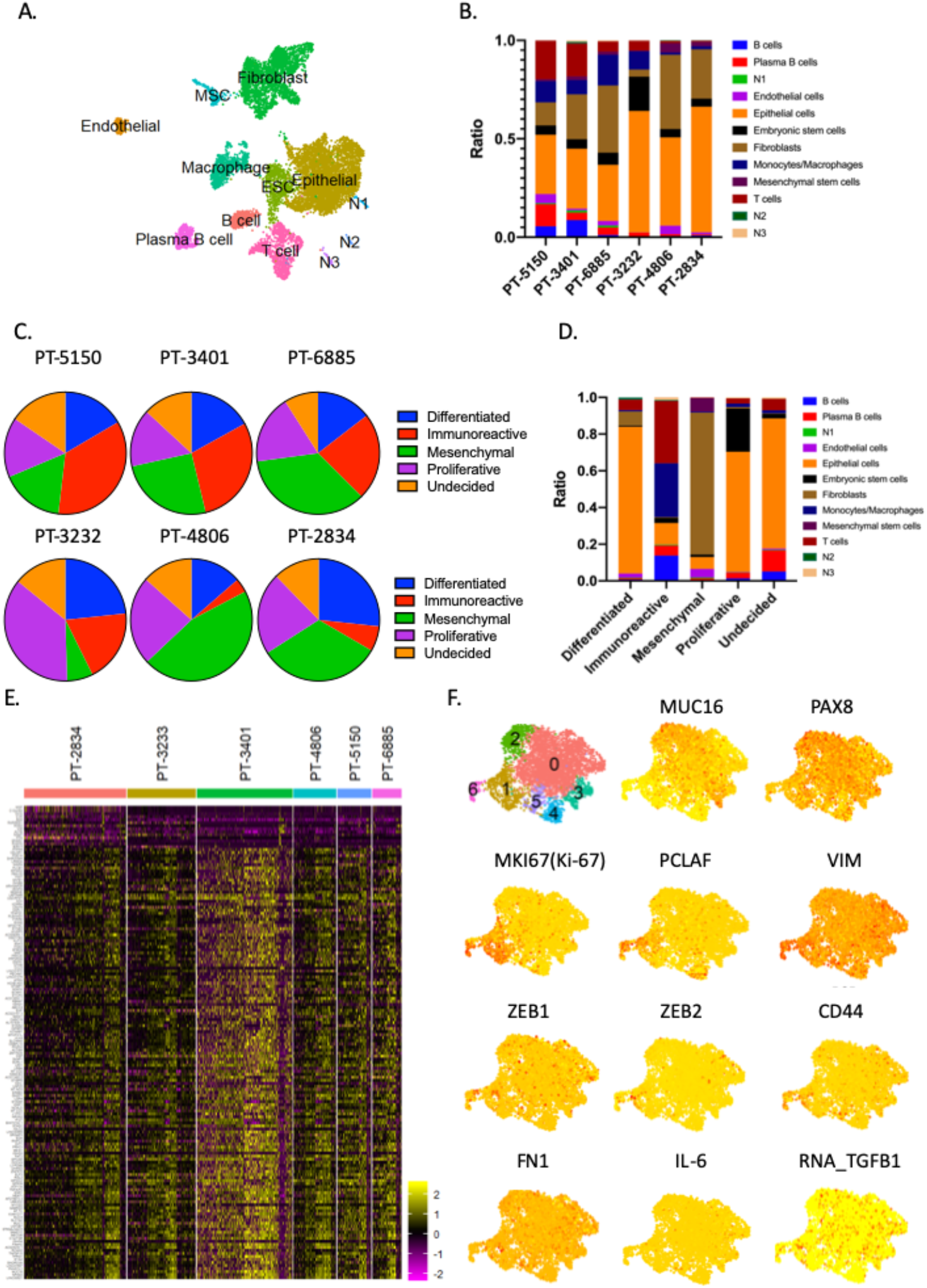
Cell and molecular subtypes assignment using Drop-seq data. A) UMAP plot for high-quality cells, based on number of genes detected per cell in all six omentum samples, colored by clustering results. B) Ratio of cellular composition by patients. C) Cancer subtype designation for each patient sample based on TCGA. D) Cancer subtype designation by cell type based on TCGA. E) Heatmap of top 150 genes for single cells (cancer and ESCs) in each patient sample. F) UMAP and feature plots of relevant marker genes of 4733 annotated cancer cell-types only from all six omentum.

It has been suggested that the TCGA subtypes may be useful in specifically categorizing cancer cells within a tumor. However, most of the cancer cells belonged to the differentiated, proliferative and undecided subtype. Therefore, to determine patient-derived differences in cancer cells that might be useful in personalizing therapy, we generated a heatmap from differentially expressed genes from each patient. The expression profile of all patients appeared similar in all patients except PT-2834 (Figure 2E). This patient was histologically diagnosed with MMMT with chondrosarcomatous elements, also confirmed by our Drop-seq data (Supplementary Figure 3). Next, the central cluster (epithelial cells) and connecting ESCs were selected and assessed for known ovarian cancer cell markers with 7 clusters detected (Figure 2F). PAX8 and MUC16, indicative of advanced disease, was expressed by multiple clusters^30,31^. Cluster 1 expressed vimentin along with high Ki-67 and PCLAF indicative of a proliferating population. Cluster 5 also expressed PCLAF, a marker of proliferation, as well as FN1 and ZEB1, markers of EMT. Cluster 6 highly expressed vimentin and Ki-67 along with comparatively lower expression of EMT markers (ZEB1 and ZEB2), suggestive of cells undergoing mesenchymal to epithelial transition (Figure 3E). Ovarian cancer stem cells marked by co-expression of CD33, CD44, CD117 and CD24 expression were not observed^32^. Cells in this cluster expressed IL-6 which correlates with poor prognosis in ovarian cancer and TGF-β which induces an aggressive cancer phenotype^33–36^. These results reveal a snapshot of the heterogeneous processes that cancer cells harness to progress disease.

**Figure 3.**
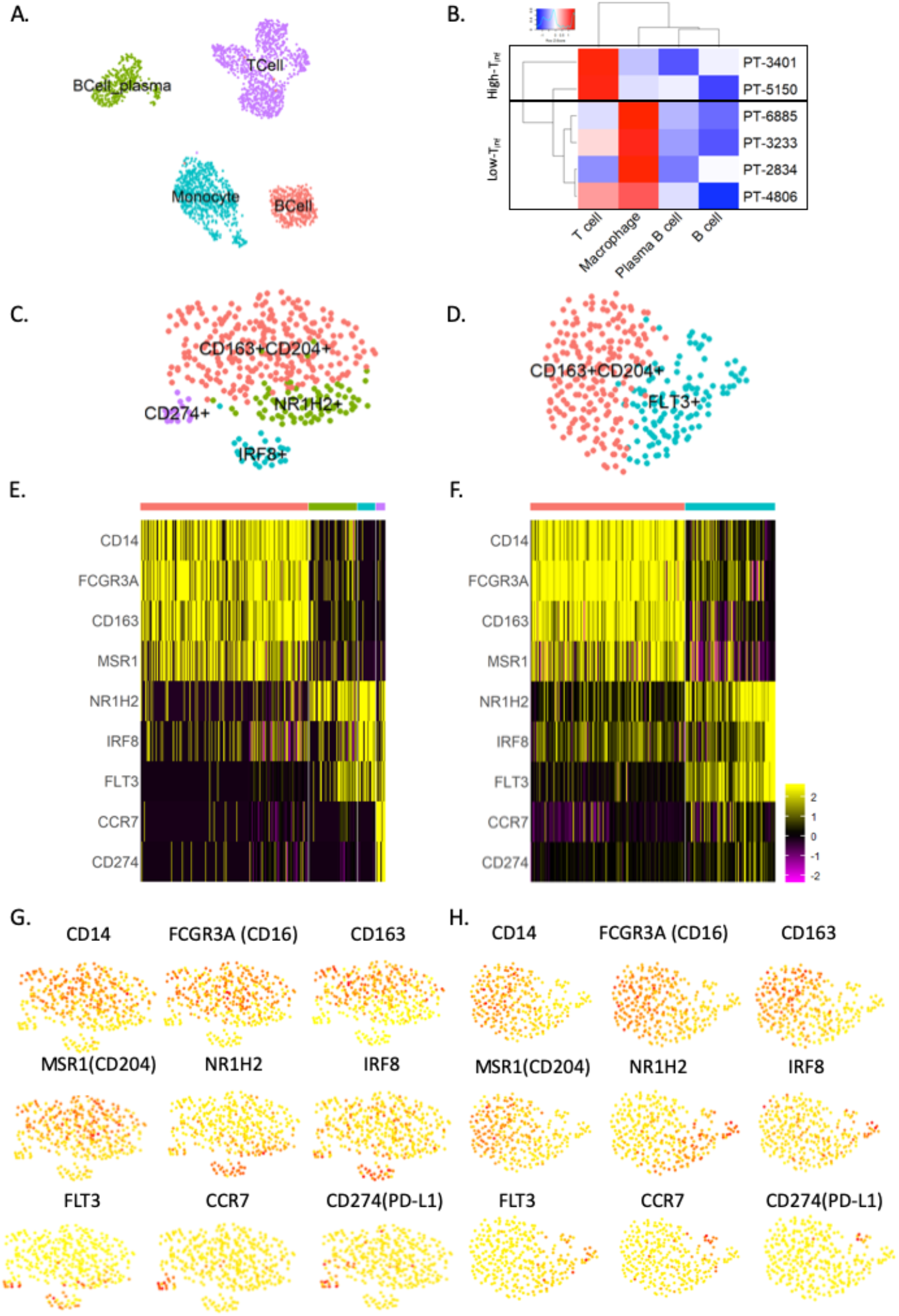
scRNA-seq of annotated immune population from omental tumors. A) UMAP plot separated by colors showing the 4 main immune cell-types based on correlation with Cell Atlas cell-type. B) Heatmap of immune cell types (T cell, B cell, plasma B cell, and Monocyte) in each patient sample with dendrograms on cell types (columns) and patients (rows), dividing the samples into high and low T cell infiltration groups. C & D) UMAP plots of unsupervised clustering of annotated macrophages from C) high T_inf_ (383 cells), and D) low T_inf_ (312 cells) groups. Heat maps from single cell analysis showing differentially expressed markers between clusters in the two groups: E) high T_inf_, and F) low T_inf_ group. G&H) Feature plots demonstrating expression of relevant genes in macrophages in high and low high T_inf_ groups, respectively.

**Figure 4.**
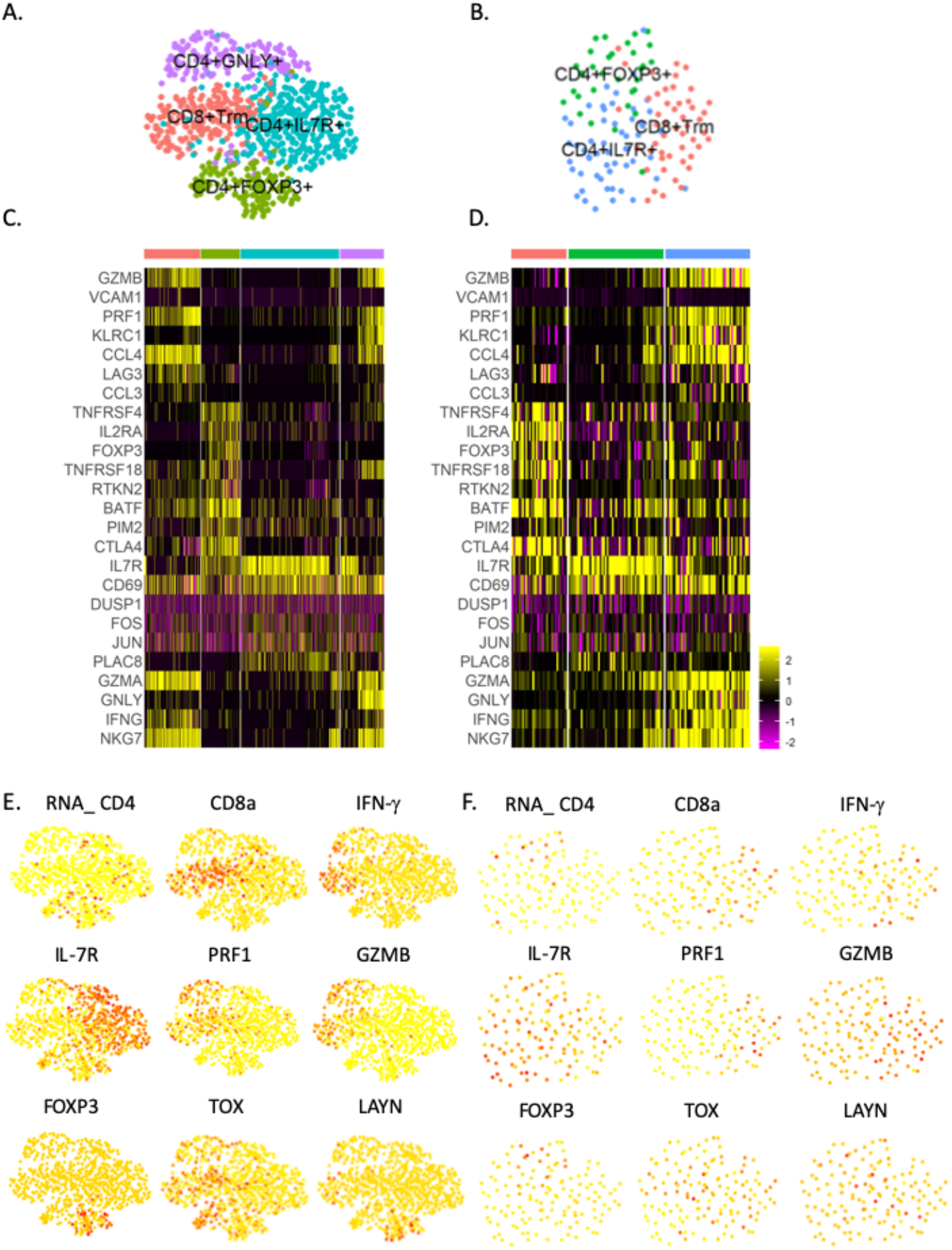
Characterization of annotated T cell population. UMAP plot for T cells in A) high T_inf_ (820 cells), and B) low T_inf_ (136) group. Heat maps from single-cell analysis of key genes in different clusters in C) high T_inf_ and D) low T_inf_ groups. Clusters in heat map are indicated by the same color as in the UMAP plots. Feature plots showing expression of key genes in E) high T_inf_, and F) low T_inf_ groups.

### Immune cellular profile of patient samples

To investigate the immune population in our patient samples, we selected the immune cells from the six patient samples and they clustered into 4 main populations; T cells, B cells, plasma B cells and monocytes/macrophages (Figure 3A). Similar samples were grouped by a dendrogram with a major difference in T cell infiltration (T_inf_) (Figure 3B). Interestingly, the high T_inf_ group (PT-5150 and PT-3401) had the lowest disease score while the low T_inf_ group (PT-4806, PT-6885, PT-2834, PT-3233) had higher disease score. During the initial phase of metastasis, macrophages are recruited from the peritoneal cavity without any anti-tumor effect on the cancer cells^37,38^. To assess the differences in the macrophage population, we performed unsupervised clustering on both groups separately (Figure 3C&D). Both groups had a CD14^+^CD16^+^CD163^+^CD204^+^ cluster reminiscent of tumor associated macrophages or TAMs (Figure 3E&F, Supplementary Figure 4). In the low T_inf_ group, there was also an immature FLT3^+^ population. These FLT3^+^ progenitors can differentiate into osteoclasts, dendritic cells, microglia and macrophages^39^. Other clusters in the high T_inf_ group included a CD274^+^ cluster, suggesting a regulatory population similar to MDSCs and an IRF8^+^ cluster induced in the presence of IFN-g and promotes the formation of autophagosome (Figure 3E&F, Supplementary Figure 4)^40^. The final cluster highly expressed NR1H2 which inhibits inflammatory genes in macrophages^41,42^. This clustering analysis reveals phenotypically distinct macrophages present in the tumor in addition to the established TAMs.

### Differences in T cell clustering and subtype analysis

To reveal the functional subtypes of T cells and any difference in the low T_inf_ versus high T_inf_ group, we clustered 136 and 820 T cells respectively (Figure 5A&B). A total of 3 and 4 transcriptionally distinct clusters emerged from low T_inf_ and high T_inf_ T cells respectively. The top genes revealed clusters similar to previously described T cell phenotypes in breast cancer, including CD4^+^IL7R^+^, CD4^+^FOXP3^+^, resident memory CD8^+^T cells (CD8^+^Trm cells) and one population described in lung cancer with high granulysin expression CD4^+^GNLY^+^ (Figure 5C-F, Supplementary Figure 5)^43,44^. Both groups had CD8^+^Trm cells which expressed IFN-g, CD4^+^IL7R^+^ and CD4^+^FOXP3^+^ T cells. The high T_inf_ group had an extra CD4^+^GNLY^+^ cluster. Contrastingly, GNLY was expressed in the low T_inf_ group by CD8^+^Trm cluster (Figure 5C&D). TOX was highly expressed by the CD8^+^Trm and the CD4^+^GNLY^+^ clusters in the high T_inf_ group as compared to the low T_inf_ group (Figure 5E&F). These TOX^+^ T cells persist during chronic infection and TOX is expressed on the CD8^+^T cells that are reactivated in response to PD-L1 immunotherapy^45,46^. Layilin, a previously described marker of exhaustion was mainly expressed by the CD4^+^FOXP3^+^ cluster (Figure 5E&F). These data suggest that scRNA-seq is capable of determining patients that will benefit from novel therapy.

**Figure 5.**
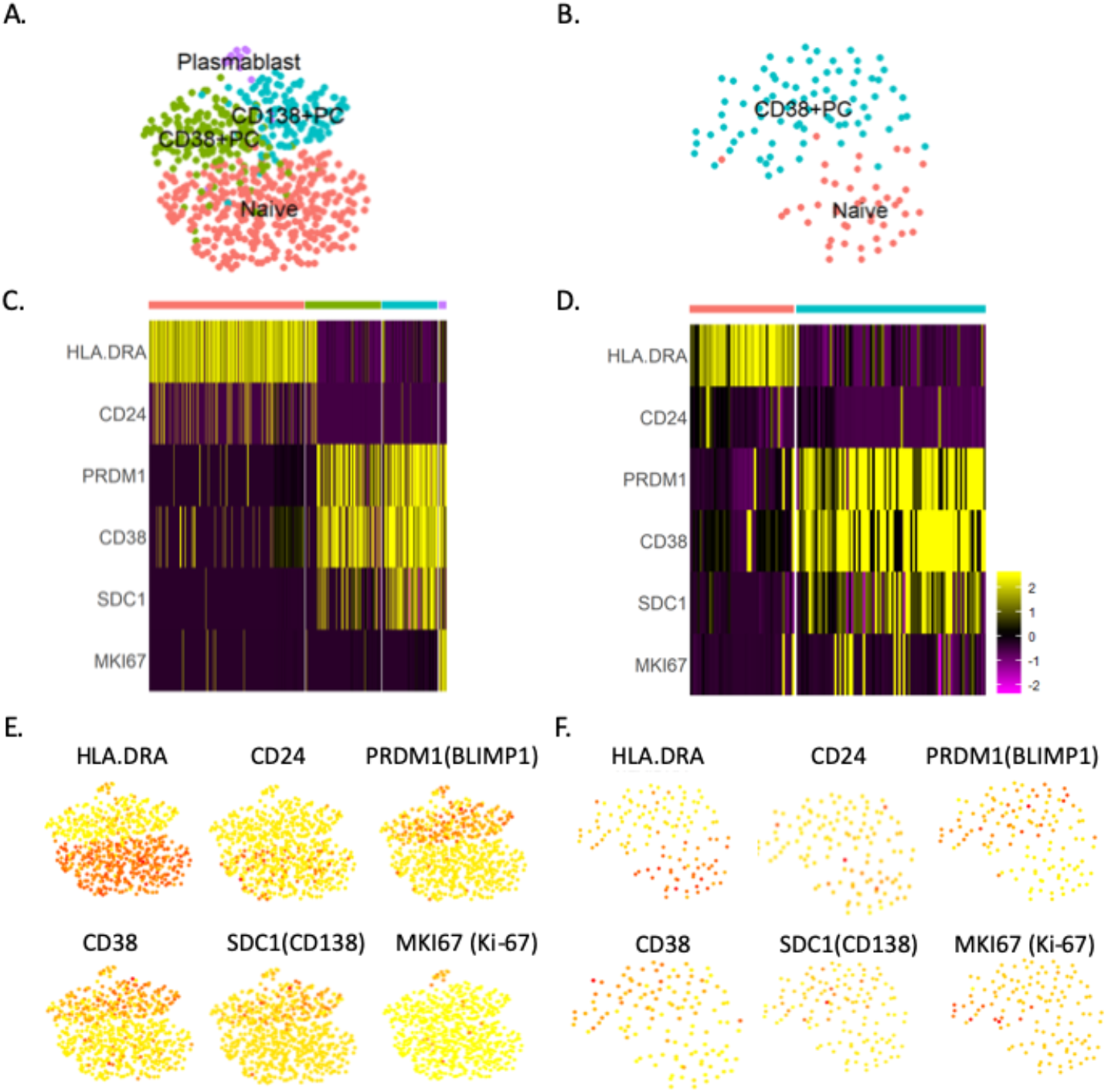
Identification of B cell clusters using single cell analysis. UMAP plot for B cells in A) high T_inf_ (396 cells) and B) low T_inf_ (124 cells) groups. Heat maps from single-cell analysis of key genes in different dusters in C) high T_inf_, and D) low T_inf_ groups. Clusters in heat map are indicated by the same color as in the UMAP plots. Feature plots showing expression of key genes in E) high T_inf_, and F) low T_inf_ groups.

### Transcriptionally distinct plasmablasts cluster in high T_inf_ group

The high T_inf_ group had an accompanying high B cell signature (Figure 3B). To investigate the B cell subtypes in our samples, naïve B cells and plasma cells were clustered yielding 4 and 2 clusters in high T_inf_ and low T_inf_ group respectively (Figure 6A&B). In the high and low T_inf_ group, the naïve cluster expressed MHC class II genes highly with a difference in CD24 expression (Figure 6C&D, Supplementary Figure 6). Other clusters in both groups expressed BLIMP1 indicating they are plasma cell clusters. In the high T_inf_ group, the BLIMP1^+^ plasma cells (PC) segregated into 3 clusters. The CD38^+^PC and CD138^+^PC are indicative of post germinal center B cells. The final cluster highly expressed MHC class II genes as well as proliferative genes suggesting a plasmablast population.

## Discussion

In this study, we characterized samples from 6 ovarian cancer patients. One to three sections of tumors were processed to account for intratumor heterogeneity based on availability. Initial IHC staining of these sections revealed a positive correlation between the area of tumor made up of adipocytes and the number of immune cells within the tumor. We processed these tumor sections and obtained Drop-seq data for a total of 9885 cells after quality control. Mapping our cells to curated cell types from the CellAtlas along with gene ontology and pathway-based enrichment was robust in assigning cells types to almost all clusters in our analysis. Our data also revealed a positive correlation between the presence of adipocytes (by IHC) and immune cells and stratified our patient cohort into two groups based on the intrinsic properties of the immune response.

The largest cluster of cells was cancer epithelial cells, which composed about 50% of cells analyzed. These cells expressed various genes associated with metastatic disease including MUC16 and PAX8, consistent with previous reports. The gene expression matrix was similar between cancer cells from each patient. The variety of cancer associated genes expressed by these cancer cells confirm the compensatory mechanisms that continue to progress disease. Another approach to suppressing tumor growth is by identifying and targeting cancer stem cell (CSC) in ovarian cancer^47^. Like other studies, we were unable to identify cells that expressed known makers of stem cells^48,49^. However, we identified a cell population closely resembling ESC and connected to the epithelial cell cluster that highly expressed proliferative marker, Ki-67. Successful identification for isolation and interrogation of putative ESC might provide useful insight about CSC in ovarian cancer.

Infiltration of immune cells into the omentum during metastatic cancer does not always elicit anti-tumor responses^38^. In ovarian cancer, high tumor infiltrating T cells is associated with significantly longer overall survival^15^. There is evidence that associates immunogenicity of neo-antigens generated due to high tumor mutational load and clinical response to immunotherapies^50–53^. To this end, we sought to characterize the differences between the two groups in our patient cohort stratified by scRNA-seq data. Macrophages make up 70% of the leukocytes present in the normal omentum so we characterized them first. Although TAMs were found in both groups, an extra 2 clusters expressing NR1H2 and CD274 were found in the high T_inf_ group. Both these markers are upregulated in an activation dependent manner suggesting that the high T_inf_ group is actually mounting an anti-tumor immune response. Flt3^+^ progenitors were unique to the low T_inf_ group. Along with the presence of TAMs, these data suggest that macrophages in the low T_inf_ group acquired immunosuppressive phenotype in response to the presence of cancer cells.

Advances in personalized medicine has revealed TOX as a transcription factor expressed by T cells that respond to immune checkpoint block^45,54^. In our patient cohort, TOX was mainly expressed by two T cell clusters, CD8^+^Trm and CD4^+^GNLY in the high T_inf_ group. The low T_inf_ group also had the CD8^+^Trm subset which expressed granulysin but not a separate CD4^+^GNLY subset. Interestingly, there was minimal TOX expression in the low T_inf_ group. Layilin, a marker gene expressed by exhausted CD8^+^T cells and regulatory T cells, was mainly expressed by the CD4^+^FOXP3^+^ cluster in our study. Only a few CD8^+^T cells expressed Layilin. Coincidentally, there is also a difference in the B cell population between the low and high T_inf_ groups. Along with the expression of TOX in T cells, the high T cell response group also had a unique subset of BLIMP1^+^CD38^+^CD138^+^ B cells and plasmablasts. It is likely that antigen-cognate plasmablasts are involved in the generation or maintenance of TOX^+^ T cells. A mechanistic study of these cells is necessary as they may hold the key into inducing TOX^+^T cells in patients who do not respond to immune checkpoint inhibitors.

In addition to the major cell types analyzed, there were other clusters including 3 undecideds (N1, N2, N3), 1 endothelial, 1 mesenchymal stem cell and 1 fibroblast cluster. For the undecided clusters, by mapping them to the CellAtlas cell types, they best correlate with astrocytes, CMP/bone marrow progenitor and plasmacytoid dendritic cells respectively. We did not recover adipocytes as they were probably lost during the tissue dissociation. Endothelial cells are the major cell types involved in angiogenesis which is necessary for metastasis. The mesenchymal stem cells are connected to the fibroblasts. Inflamed omentum contains stem cells displaying similar surface markers to MSCs. These stem cells are capable of differentiating into fat, cartilage or bone depending on the secreted factors present^55,56^. Finally, there was a significant fibroblast cluster in our sample population. Metastatic transformation of the omentum changes the cellular composition from mainly adipocytes to cancer cells, immune cells and fibroblasts^28^. Cancer associated fibroblasts mainly function to remodel the extracellular matrix in the tumor microenivironment^57^. Recently, a scRNA-seq study of CAF in pancreatic cancer revealed a LRRC15^+^ CAF population that correlated with poor response in patients treated with anti-PD-L1 therapy^58^. A few of the fibroblasts in our study also express LRRC15 (data not shown). Ultimately, more studies are needed to further elucidate these cells and how they can be manipulated to enhance immunotherapeutic approaches.

## Conclusion

To our knowledge, this is the first scRNA-seq study that allows stratification of patient samples all initially pathologically classified as stage III, according to immune response in metastatic ovarian cancer. Concurrent transcriptomic analysis of cancer and stromal cells revealed patient heterogeneity highlighting the need for personalized medicine. Interrogating tumor infiltrating lymphocytes at the single cell level also revealed patient-intrinsic responses that are previously undescribed in ovarian cancer. Mechanistic studies are required to determine the role of these cells in tumor progression and response to novel cancer immunotherapies.

## Acknowledgment

We are grateful to Drs. Mengjie Chen and Ernst Lengyel for helpful discussions and Mark Lingen for tissue access. All tissues were histologically graded, de-identified and obtained from the University of Chicago Human Tissue Resource Center. Computational resources and data storage were provided by the University of Chicago Research Computing Center. This work was performed, in part, at the Center for Nanoscale Materials, a U.S. Department of Energy Office of Science User Facility, and supported by the U.S. Department of Energy, Office of Science, under Contract No. DE-AC02-06CH11357.

## Conflict of interest

The authors declare no potential conflict of interest.

## Funding

This project is internally funded.

## References

1 Siegel, R. L., Miller, K. D. & Jemal, A. Cancer Statistics, 2017. CA: a cancer journal for clinicians 67, 7–30, doi:10.3322/caac.21387 (2017).

2 Testa, U., Castelli, G. & Pelosi, E. Genetic Abnormalities, Clonal Evolution, and Cancer Stem Cells of Brain Tumors. Med Sci (Basel) 6, doi:10.3390/medsci6040085 (2018).

3 Tothill, R. W. et al. Novel molecular subtypes of serous and endometrioid ovarian cancer linked to clinical outcome. Clinical cancer research: an official journal of the American Association for Cancer Research 14, 5198–5208, doi:10.1158/1078-0432.CCR-08-0196 (2008).

4 Network, C. G. A. R. Integrated genomic analyses of ovarian carcinoma. Nature 474, 609 (2011).

5 Zheng, G. X. et al. Massively parallel digital transcriptional profiling of single cells. Nat Commun 8, 14049, doi:10.1038/ncomms14049 (2017).

6 Villani, A. C. et al. Single-cell RNA-seq reveals new types of human blood dendritic cells, monocytes, and progenitors. Science 356, doi:10.1126/science.aah4573 (2017).

7 Dixit, A. et al. Perturb-Seq: Dissecting Molecular Circuits with Scalable Single-Cell RNA Profiling of Pooled Genetic Screens. Cell 167, 1853–1866 e1817, doi:10.1016/j.cell.2016.11.038 (2016).

8 Jaitin, D. A. et al. Dissecting Immune Circuits by Linking CRISPR-Pooled Screens with Single-Cell RNA-Seq. Cell 167, 1883–1896 e1815, doi:10.1016/j.cell.2016.11.039 (2016).

9 Shalek, A. K. et al. Single-cell transcriptomics reveals bimodality in expression and splicing in immune cells. Nature 498, 236–240, doi:10.1038/nature12172 (2013).

10 Zheng, C. et al. Landscape of Infiltrating T Cells in Liver Cancer Revealed by Single-Cell Sequencing. Cell 169, 1342–1356 e1316, doi:10.1016/j.cell.2017.05.035 (2017).

11 Hu, Z. et al. The Repertoire of Serous Ovarian Cancer Non-genetic Heterogeneity Revealed by Single-Cell Sequencing of Normal Fallopian Tube Epithelial Cells. Cancer cell 37, 226–242 e227, doi:10.1016/j.ccell.2020.01.003 (2020).

12 Platell, C., Cooper, D., Papadimitriou, J. M. & Hall, J. C. The omentum. World J Gastroenterol 6, 169–176, doi:10.3748/wjg.v6.i2.169 (2000).

13 Clark, R. et al. Milky spots promote ovarian cancer metastatic colonization of peritoneal adipose in experimental models. The American journal of pathology 183, 576–591, doi:10.1016/j.ajpath.2013.04.023 (2013).

14 Nieman, K. M. et al. Adipocytes promote ovarian cancer metastasis and provide energy for rapid tumor growth. Nature medicine 17, 1498–1503, doi:10.1038/nm.2492 (2011).

15 Santoiemma, P. P. & Powell, D. J., Jr. Tumor infiltrating lymphocytes in ovarian cancer. Cancer Biol Ther 16, 807–820, doi:10.1080/15384047.2015.1040960 (2015).

16 Macosko, E. Z. et al. Highly Parallel Genome-wide Expression Profiling of Individual Cells Using Nanoliter Droplets. Cell 161, 1202–1214, doi:10.1016/j.cell.2015.05.002 (2015).

17 Frankish, A. et al. GENCODE reference annotation for the human and mouse genomes. Nucleic Acids Res 47, D766–D773, doi:10.1093/nar/gky955 (2019).

18 Dobin, A. et al. STAR: ultrafast universal RNA-seq aligner. Bioinformatics 29, 15–21, doi:10.1093/bioinformatics/bts635 (2013).

19 Selewa, A. et al. Systematic Comparison of High-throughput Single-Cell and Single-Nucleus Transcriptomes during Cardiomyocyte Differentiation. BioRxiv, doi:10.1101/585901 (2019).

20 Butler, A., Hoffman, P., Smibert, P., Papalexi, E. & Satija, R. Integrating single-cell transcriptomic data across different conditions, technologies, and species. Nature biotechnology 36, 411–420, doi:10.1038/nbt.4096 (2018).

21 Stuart, T. et al. Comprehensive Integration of Single-Cell Data. Cell 177, 1888–1902 e1821, doi:10.1016/j.cell.2019.05.031 (2019).

22 Stuart, T. et al. Comprehensive Integration of Single-Cell Data. Cell 177, 1888–1902 (2019).

23 McInnes, L., Healy, J. & Melville, J. UMAP: Uniform Manifold Approximation and Projection for Dimension Reduction. Journal of Open Source Software 29, 861, doi:https://doi.org/10.21105/joss.00861 (2018).

24 Maaten, L. v. d. & Hinton, G. isualizing data using t-SNE. Journal of machine learning research 9 9, 2579–2605 (2008).

25 I, T. et al. Dissecting the multicellular ecosystem of metastatic melanoma by single-cell RNA-seq. Science 352, 189–196 (2016).

26 Mabbott, N. A., Baillie, J. K., Brown, H., Freeman, T. C. & Hume, D. A. An expression atlas of human primary cells: inference of gene function from coexpression networks. BMC Genomics 14, 632, doi:10.1186/1471-2164-14-632 (2013).

27 Waltman, L. & Eck, N. J. V. A smart local moving algorithm for large-scale modularity-based community detection. The European physical journal B 86, 471 (2013).

28 Pearce, O. M. T. et al. Deconstruction of a Metastatic Tumor Microenvironment Reveals a Common Matrix Response in Human Cancers. Cancer Discov 8, 304–319, doi:10.1158/2159-8290.CD-17-0284 (2018).

29 Network., C. G. A. R. Intergrated genomic analyses of ovarian carcinoma. Nature 474, 609–615, doi:https://doi.org/10.1038/nature10166 (2011).

30 Robertson, D. M. et al. Combined inhibin and CA125 assays in the detection of ovarian cancer. Clin Chem 45, 651–658 (1999).

31 Theriault, C. et al. MUC16 (CA125) regulates epithelial ovarian cancer cell growth, tumorigenesis and metastasis. Gynecologic oncology 121, 434–443, doi:10.1016/j.ygyno.2011.02.020 (2011).

32 Klemba, A. et al. Surface markers of cancer stem-like cells of ovarian cancer and their clinical relevance. Contemp Oncol (Pozn) 22, 48–55, doi:10.5114/wo.2018.73885 (2018).

33 Coward, J. et al. Interleukin-6 as a therapeutic target in human ovarian cancer. Clinical cancer research: an official journal of the American Association for Cancer Research 17, 6083–6096, doi:10.1158/1078-0432.CCR-11-0945 (2011).

34 Lutgendorf, S. K. et al. Interleukin-6, cortisol, and depressive symptoms in ovarian cancer patients. Journal of clinical oncology: official journal of the American Society of Clinical Oncology 26, 4820–4827, doi:10.1200/JCO.2007.14.1978 (2008).

35 Scambia, G. et al. Prognostic significance of interleukin 6 serum levels in patients with ovarian cancer. Br J Cancer 71, 354–356, doi:10.1038/bjc.1995.71 (1995).

36 Li, W. et al. TGFbeta1 in fibroblasts-derived exosomes promotes epithelial-mesenchymal transition of ovarian cancer cells. Oncotarget 8, 96035–96047, doi:10.18632/oncotarget.21635 (2017).

37 Shimotsuma, M., Simpson-Morgan, M. W., Takahashi, T. & Hagiwara, A. Activation of omental milky spots and milky spot macrophages by intraperitoneal administration of a streptococcal preparation, OK-432. Cancer research 52, 5400–5402 (1992).

38 Oosterling, S. J. et al. Insufficient ability of omental milky spots to prevent peritoneal tumor outgrowth supports omentectomy in minimal residual disease. Cancer Immunol Immunother 55, 1043–1051, doi:10.1007/s00262-005-0101-y (2006).

39 Servet-Delprat, C. et al. Flt3+ macrophage precursors commit sequentially to osteoclasts, dendritic cells and microglia. BMC Immunol 3, 15, doi:10.1186/1471-2172-3-15 (2002).

40 Gupta, M. et al. IRF8 directs stress-induced autophagy in macrophages and promotes clearance of Listeria monocytogenes. Nat Commun 6, 6379, doi:10.1038/ncomms7379 (2015).

41 Castrillo, A., Joseph, S. B., Marathe, C., Mangelsdorf, D. J. & Tontonoz, P. Liver X receptor-dependent repression of matrix metalloproteinase-9 expression in macrophages. J Biol Chem 278, 10443–10449, doi:10.1074/jbc.M213071200 (2003).

42 N, A. G. & Castrillo, A. Liver X receptors as regulators of macrophage inflammatory and metabolic pathways. Biochim Biophys Acta 1812, 982–994, doi:10.1016/j.bbadis.2010.12.015 (2011).

43 Savas, P. et al. Publisher Correction: Single-cell profiling of breast cancer T cells reveals a tissue-resident memory subset associated with improved prognosis. Nature medicine 24, 1941, doi:10.1038/s41591-018-0176-6 (2018).

44 Guo, X. et al. Publisher Correction: Global characterization of T cells in non-small-cell lung cancer by single-cell sequencing. Nature medicine 24, 1628, doi:10.1038/s41591-018-0167-7 (2018).

45 Yao, C. et al. Single-cell RNA-seq reveals TOX as a key regulator of CD8(+) T cell persistence in chronic infection. Nat Immunol 20, 890–901, doi:10.1038/s41590-019-0403-4 (2019).

46 Khan, O. et al. TOX transcriptionally and epigenetically programs CD8(+) T cell exhaustion. Nature 571, 211–218, doi:10.1038/s41586-019-1325-x (2019).

47 Bast, R. C., Jr., Hennessy, B. & Mills, G. B. The biology of ovarian cancer: new opportunities for translation. Nat Rev Cancer 9, 415–428, doi:10.1038/nrc2644 (2009).

48 Shah, M. M. & Landen, C. N. Ovarian cancer stem cells: are they real and why are they important? Gynecologic oncology 132, 483–489, doi:10.1016/j.ygyno.2013.12.001 (2014).

49 Burgos-Ojeda, D., Rueda, B. R. & Buckanovich, R. J. Ovarian cancer stem cell markers: prognostic and therapeutic implications. Cancer letters 322, 1–7, doi:10.1016/j.canlet.2012.02.002 (2012).

50 Van Allen, E. M. et al. Genomic correlates of response to CTLA-4 blockade in metastatic melanoma. Science 350, 207–211, doi:10.1126/science.aad0095 (2015).

51 Rizvi, N. A. et al. Cancer immunology. Mutational landscape determines sensitivity to PD-1 blockade in non-small cell lung cancer. Science 348, 124–128, doi:10.1126/science.aaa1348 (2015).

52 Snyder, A. et al. Genetic basis for clinical response to CTLA-4 blockade in melanoma. N Engl J Med 371, 2189–2199, doi:10.1056/NEJMoa1406498 (2014).

53 Aguade-Gorgorio, G. & Sole, R. Genetic instability as a driver for immune surveillance. J Immunother Cancer 7, 345, doi:10.1186/s40425-019-0795-6 (2019).

54 Siddiqui, I. et al. Intratumoral Tcf1(+)PD-1(+)CD8(+) T Cells with Stem-like Properties Promote Tumor Control in Response to Vaccination and Checkpoint Blockade Immunotherapy. Immunity 50, 195–211 e110, doi:10.1016/j.immuni.2018.12.021 (2019).

55 Shah, S. et al. Cellular basis of tissue regeneration by omentum. PloS one 7, e38368, doi:10.1371/journal.pone.0038368 (2012).

56 Friedenstein, A. J., Petrakova, K. V., Kurolesova, A. I. & Frolova, G. P. Heterotopic of bone marrow. Analysis of precursor cells for osteogenic and hematopoietic tissues. Transplantation 6, 230–247 (1968).

57 Kalluri, R. The biology and function of fibroblasts in cancer. Nat Rev Cancer 16, 582–598, doi:10.1038/nrc.2016.73 (2016).

58 Dominguez, C. X. et al. Single-Cell RNA Sequencing Reveals Stromal Evolution into LRRC15(+) Myofibroblasts as a Determinant of Patient Response to Cancer Immunotherapy. Cancer Discov 10, 232–253, doi:10.1158/2159-8290.CD-19-0644 (2020).

