## Supplementary figures for "Characterizing the tumor microenvironment of metastatic ovarian cancer by single cell transcriptomics"

| Cell-type | Min. | Median | Mean |
|---|---|---|---|
| All | 601 | 1328 | 1742 |
| Cancer | 604 | 1622 | 2105 |
| Immune | 601 | 976 | 1098 |
| Other | 608 | 1449 | 1734 |
| B Cell | 602 | 901 | 950 |
| Plasma B Cell | 606 | 1123 | 1250 |
| Dendritic Cell Plasmacytoid | 677 | 1059 | 1209 |
| Endothelial | 608 | 1189 | 1599 |
| Epithelial | 604 | 1576 | 2044 |
| ESC | 632 | 2007 | 2518 |
| Fibroblast | 618 | 1472 | 1730 |
| Macrophages | 601 | 1127 | 1269 |
| MSC | 666 | 1255 | 1574 |
| T cell | 602 | 923 | 973 |

Supplementary table 1. Table for number of genes detected per cell, broken down by annotated cell types.

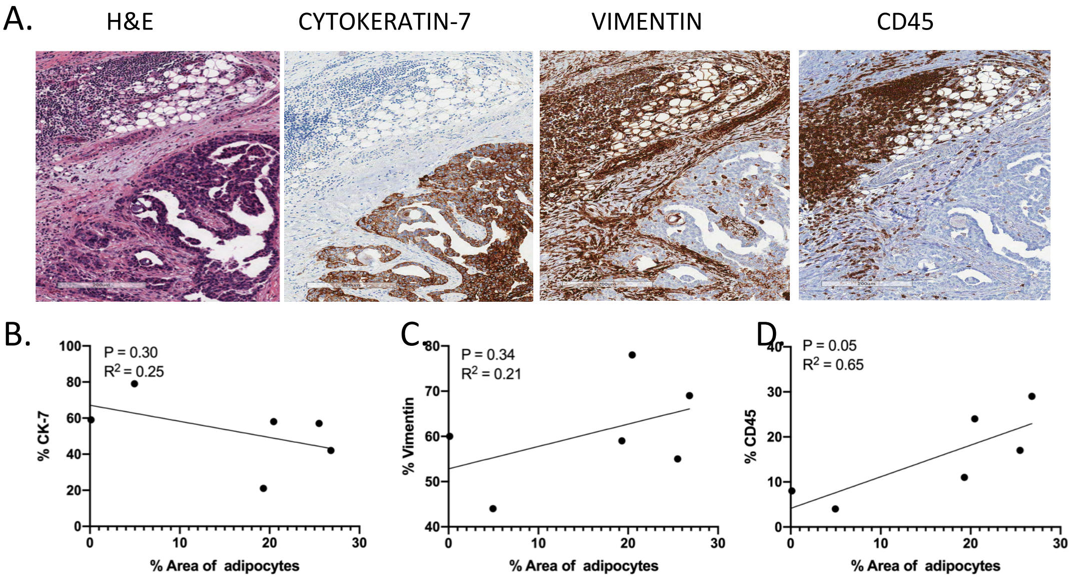


Supplementary Figure 1. A) Images of H&E and IHC staining of one sample shown at 400x. B) One way ANOVA calculated between area of adipocytes and B) %CK-7, C) % vimentin positive cells, and D) % CD45 positive immune cells.

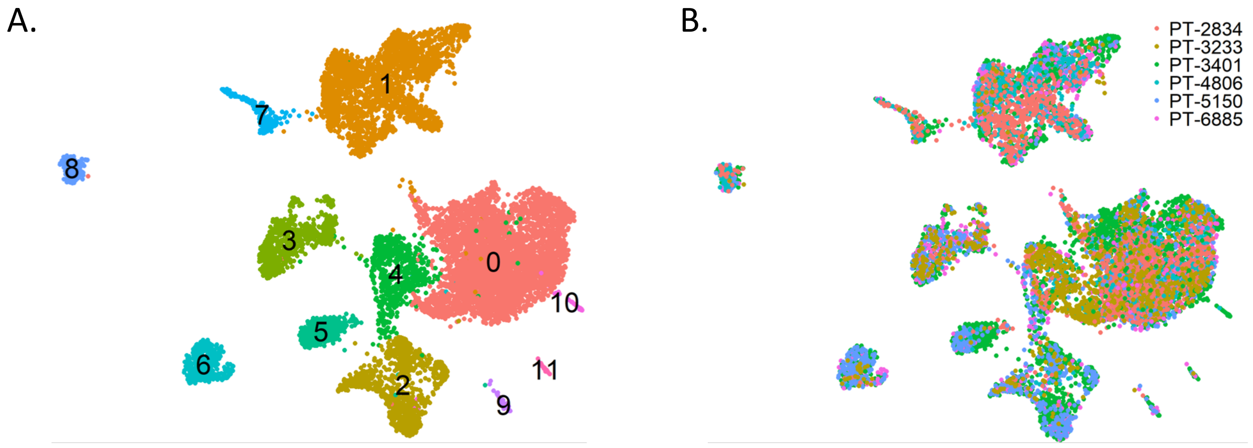


Supplementary Figure 2. A) Unsupervised clustering of 9885 single cells isolated from omental tumors from 6 patients using UMAP yield 12 clusters. B) Cell clusters shown by patient origin.

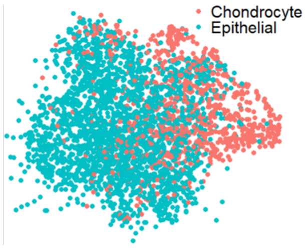


Supplementary Figure 3. UMAP of cluster for cancer cells split into epithelial and sarcoma origin.

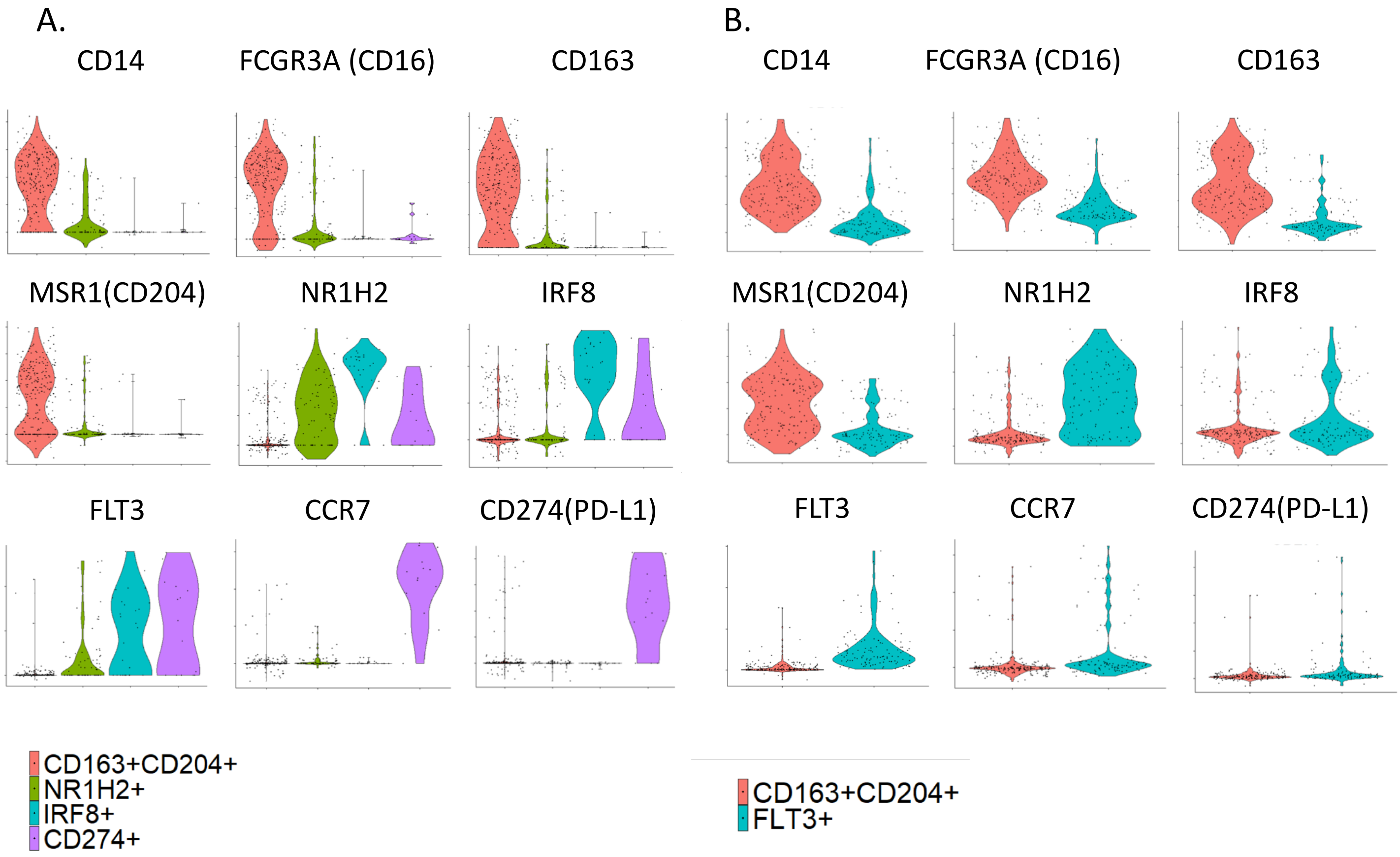


Figure 4 Violin plots of Macrophage relevant genes by cluster in A) high $T_{inf}$, and B) low $T_{inf}$ groups.

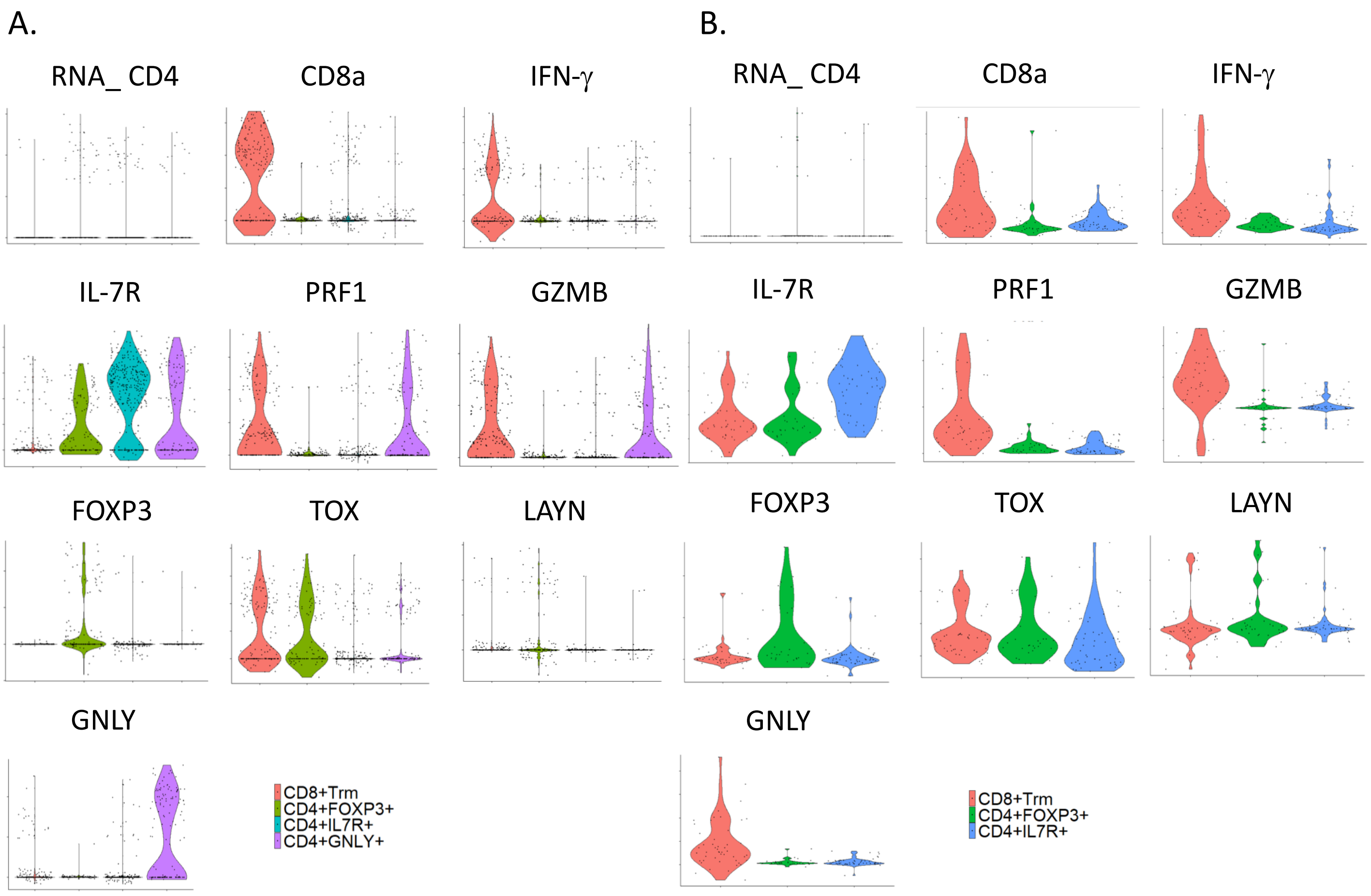


Figure 5. Violin plots of T cell relevant genes by cluster in A) high $T_{inf}$, and B) low $T_{inf}$ groups.

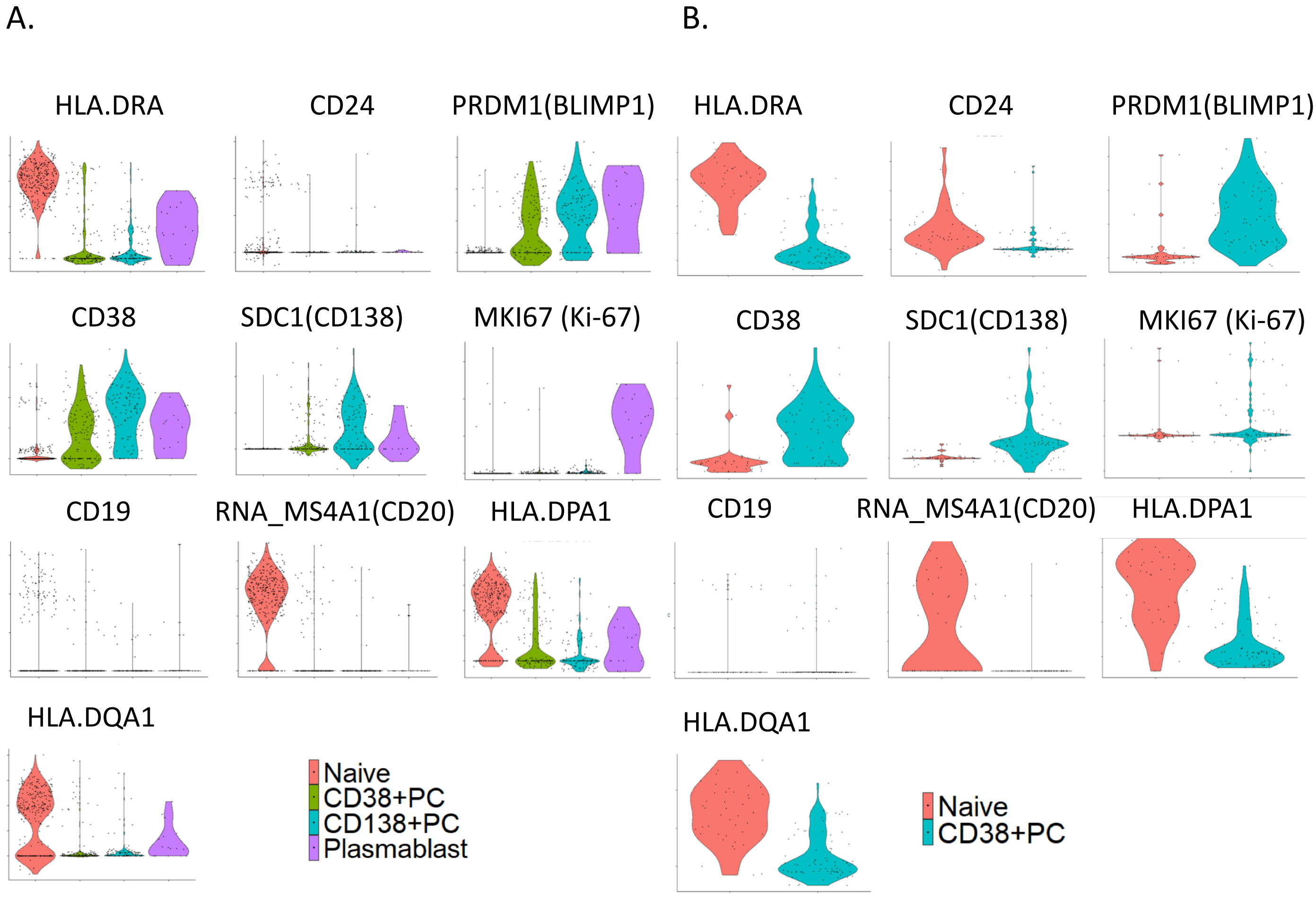


Figure 6. Violin plots of B cell relevant genes by cluster in A) high $T_{inf}$, and B) low $T_{inf}$ groups.
